# Laboratory rearing alters activity and sleep patterns in the olive fruit fly

**DOI:** 10.1101/2022.08.31.506115

**Authors:** Anastasia M. Terzidou, Dimitrios S. Koveos, Nikos T. Papadopoulos, James R. Carey, Nikos A. Kouloussis

## Abstract

Olive fruit flies, *Bactrocera oleae* (Diptera: Tephritidae) that are laboratory reared in artificial diet are essential for the genetic control techniques for this pest. However, the colony’s laboratory adaptation can affect their quality. We used the Locomotor Activity Monitor (LAM25, Trikinetics, MA, USA) to track the activity and sleep patterns of adult olive fruit flies reared as immatures in olives (F2-F3 generation) and in artificial diet (>300 generations). Counts of beam breaks caused by the adult fly activity were used as an estimation of its locomotor activity levels during the light and dark period. A longer than five minutes period of adults’ inactivity during the dark period was considered as one sleep episode. Activity levels and sleep parameters were found to be dependent on sex, mating status and rearing history. In virgin flies reared in olives, males were more active than females and increased their locomotor activity towards the end of the light period. Mating decreased the locomotor activity levels of males, but not of females olive-reared flies. Laboratory flies reared in artificial diet had lower locomotor activity levels during the light period and more sleep episodes of shorter duration compared to flies reared in olives. We describe the diurnal locomotor activity patterns of *B. oleae* adults reared in olive fruits and in artificial diet. We discuss how locomotor activity and sleep pattern differences may affect the laboratory flies’ ability to compete with wild males in the field.

## Introduction

The olive fruit fly has been the most important pest of olives in Mediterranean countries for at least 2000 years [1] and in 1998 it was first detected in California [2]. Its distribution now covers the Mediterranean basin, North and Sub-Saharan Africa, south-west Asia and North America [3]. Because of the significant economic damage caused by *B. oleae* and the intense use of insecticides to manage it, Sterile Insect Technique (SIT), as well as a self-limiting olive fly technology have been proposed to control this pest [4,5]. Both methods require the mass-release of quality laboratory male olive fruit flies reared in artificial diet, that will be able to survive, search, find, and mate effectively with the wild population. Selection during colonization and mass-rearing normally changes the biology and behavior of an insect species. A common problem in mass-reared insects is the loss of irritability, because crowded rearing conditions select for individuals that ignore movements in their surroundings [6]. The mass-rearing conditions reduce the flight capacity of *Amyelois transitella* [7], and the overall activity of *Bactrocera tryoni* compared to wild population [8], while inbreeding in *Drosophila melanogaster* decreases locomotor activity and changes daily activity patterns [9]. It has been shown that in *Bactrocera dorsalis*, the time of mating - which is controlled by the circadian clock - can be altered in insects that have been adapted in laboratory rearing conditions, even after a period of just a year [10].

Sleep and sleep-like states are present in all animals and many studies have shown sleep-like states exist even in arthropods and nematodes [11]. The circadian system is responsible for controlling daily activity patterns, such as locomotion, mating, and also sleep [12,13]. Sleep is necessary for an animal’s survival and health, including replenishment of energy stores, removal of harmful by-products, and maintenance of neural plasticity [14]. In *D. melanogaster* sleep deprivation can result to death [15, 16]. There is also evidence that night sleep disruption, including sleep deprivation (total sleep loss) and sleep restriction (partial sleep loss), caused by a variety of biotic and abiotic factors in nature can cause changes in insect daily activity patterns with consequences in behavioral performance during active periods [17]. Daily rhythms of activity and rest can be recorded by placing individuals in glass tubes and monitoring the movements using infrared beam-based activity monitors like Locomotor Activity Monitor-LAM by Trikinetics (https://trikinetics.com/) [18]. *D. melanogaster* has been used extensively as a model organism for sleep studies [19, 20] where the duration of sleep episodes or bouts during the night indicates how well a fly can stay asleep [21]. In drosophila, the 5 min threshold for defining sleep state is widely accepted [22]. One limitation of the LAM tracking system is that it cannot differentiate between inactivity and sleep [23]. For bumblebees, bouts of inactivity lasting more than 5 min during the night were visually associated with posture indicating sleep, but during the day, inactivity periods could not reliable be associated with sleep [24]. For this study, we accepted that a bout of more than 5 min of inactivity during the dark period or light period is considered a sleep or rest episode respectively.

The aim of this study was to record the diurnal patterns of locomotion of olive fruit flies, as a detailed tracking of their activity with the use of LAM. We focused on reproductively mature wild flies (F2-F3 generation reared in olive fruits) and how their diurnal patterns of locomotion change according to sex and mating status. We also studied the effect of laboratory rearing on artificial diet to the locomotor activity and sleep patterns of reproductively mature virgin male flies. We hypothesized that wild flies reared on olives would have higher locomotor activity levels than laboratory flies reared on artificial diet, as laboratory mass rearing is known to further decrease the tendency of flies to move [8].

## 2. Materials and methods

### 2.1. Insect rearing

The wild olive fruit fly colony was established with flies that emerged from infested olives field collected in late September in the area of Thessaloniki. Emerged adults were maintained in colony wooden cages (30 × 30 × 30 cm) under laboratory conditions (24±1.5 °C, RH 40±5 %, L:D 14:10), fed with a diet consisting of sugar, yeast hydrolysate and water (ratio 4:1:5) and allowed to oviposit in olives. Water was provided with a soaked cotton stick extruding from a small water container. Flies of this wild population that were grown in their larval stages in olives for 2-3 generations in our laboratory were used in our experiments (referred to as W flies hereafter).

We used laboratory adapted olive fruit flies reared in their larval stages in artificial diet (referred to as AR flies hereafter) from the colony maintained in our laboratory for more than 20 generations. The colony was established from the “Democritus strain”, which was developed at the Democritus Nuclear Research Center, Athens, Greece and had already been reared for more than 300 generations. Adult flies were kept in wooden cages (30 × 30 × 30 cm) and each cage contained about 200 individuals. Adult food was given in the form of a liquid diet consisting of sugar, yeast hydrolysate and water (ratio 4:1:5) (no antibiotic was added). Egg yolk powder was added ad libitum as extra protein source for the colony AR flies. They were allowed to oviposit on beeswax domes (diameter = 2 cm) and eggs were collected every two days with a fine brush and washed with propionic acid solution (0.3 %). The collected eggs were then placed directly on the larval diet inside a Petri dish (94 × 16 mm). Larval artificial diet consisted of 550 ml of tap water, olive oil (20 ml), Tween 80 (7.5 ml), potassium sorbate (0.5 g), nipagin (2 g), crystalline sugar (20 g), brewer’s yeast (75 g), soy hydrolysate (30 g), hydrochloric acid 2N (30 ml), and cellulose powder as bulking medium (275 g) as described in Tsitsipis et al [25]. The diet was kept moist to stimulate last stage larvae to exit the diet which were then collected by sieving the sand on which the Petri dish was placed.

Newly emerged W and AR flies were separated by sex in the first 24h of their emergence, kept in plexiglass cages (15 × 15 × 15 cm) that contained 20 flies each and under the same conditions (T: 24±1.5°C, RH: 40±5%, L:D 14:10) and fed with the same diet consisting of sugar, yeast hydrolysate and water (ratio 4:1:5).

Male and female W flies of the same cohort were kept together after adult emergence in colony wooden cages (about 70-90 flies per cage) with the same diet and environmental conditions as above. They were considered mated by the time they were used for the bioassays (12-13 days old at the beginning of bioassay).

Laboratory adaptation results in more rapid sexual maturity rate for AR flies [26, 27]. Sexual maturity begins on the 8th day of age of W flies of both sexes [28], while for AR males, it begins on the 2nd day and for AR females on the 3rd day of age. W flies used were 12-13 days old at the beginning of the bioassay, while AR flies were 6-7 days old, considering the AR flies’ shorter longevity [29] and that AR flies for SIT purposes are released at a young age [30].

### 2.2. Locomotor Activity

We recorded the locomotor activity patterns of adult olive fruit flies by using the Locomotor Activity Monitor-LAM25 system (Trikinetics). In this system flies were individually kept in each of 32 glass tubes with 25 mm diameter and 125 mm length. On the one end of the tube, we adjusted a vinyl plastic stopper (CAP25-BLK-Vinyl Tube Cap-Trikinetics) inside which was an agar-based gel diet with sugar, yeast hydrolysate, agar, nipagin and water (4:1:0.2:0.1:20) for food and water provision [31]. The other end of the tube was covered with a piece of organdie to allow ventilation. The tubes were maintained in a climatic room under a photoperiod of 14:10 L:D. The light period was from 07:00 to 21:00 (hereafter referred to as LP), and dark period was from 21:00 to 07:00 (hereafter referred to as DP). Activity at each tube was measured every minute as counts of infrared light beams crossed.

Three LAM devices were used simultaneously for the W flies bioassay. Thirty-two W virgin males and thirty-two W virgin females were maintained in two LAMs. Sixteen W mated males and sixteen W mated females were maintained in the third LAM. They were monitored for 5 consecutive days. After the completion of this bioassay, AR virgin flies (sixteen males and sixteen females) were maintained in one LAM device and monitored for 5 consecutive days.

### 2.3. Data analysis

The LAM devices were set to record the sum of movements each fly performed every minute and exported the data in monitor files as the number of counts for each tube. The raw monitor data were processed in the DAM FileScan software and activity data collected in 1-minute intervals (1-minute bins) were compressed and converted to 30-minute intervals (30-minute bins) for plotting purposes. Activity/sleep analysis was performed using an in-house MATLAB program called Sleep and Circadian Analysis MATLAB Program (SCAMP) [32]. Activity levels during the LP and DP were the total counts for each fly during the 14 h period of lights on and 10 h period of lights off respectively. We conventionally refer to bouts of > 5 min of inactivity as rest and sleep episodes when they occur during the LP and DP respectively. Activity levels, the number and duration of rest/sleep episodes were calculated as the average of 5 consecutive days of monitoring after excluding the flies that died during the bioassay. Sample size was *n* = 32 W virgin flies of each sex, *n* = 16 AR virgin flies of each sex, and *n* = 12 male and *n* = 14 female W mated flies. For all the comparisons, a 2-tailed *t-*test was performed (level of significance α = 0.05). with the statistical software package JMP 14.1.0 [33].

## 3. Results

### 3.1. Locomotor activity

#### 3.1.1. Locomotor activity of W flies

The pattern of locomotor activity of W virgin males and females during 24-h in 30 min bins is shown in Fig 1A. The mean locomotor activity level (SE) of W virgin males was during the LP was 2031.3 (132.0) counts and that of W virgin females was 1394.2 (86.6) counts. They differed significantly (2-tailed *t*-test = 4.035, *df* = 54, *P* = .0002). However, during the DP, the mean locomotor activity level (SE) of W virgin males and W virgin females was 257.2 (17.8) counts and 246.6 (16.3) counts respectively and did not differ significantly (2-tailed *t*-test = .438, *df* = 62, *P* = .662). High levels of locomotor activity during the DP were recorded at the time of lights off and for the next hour after the transition to scotophase.

**Fig 1.**
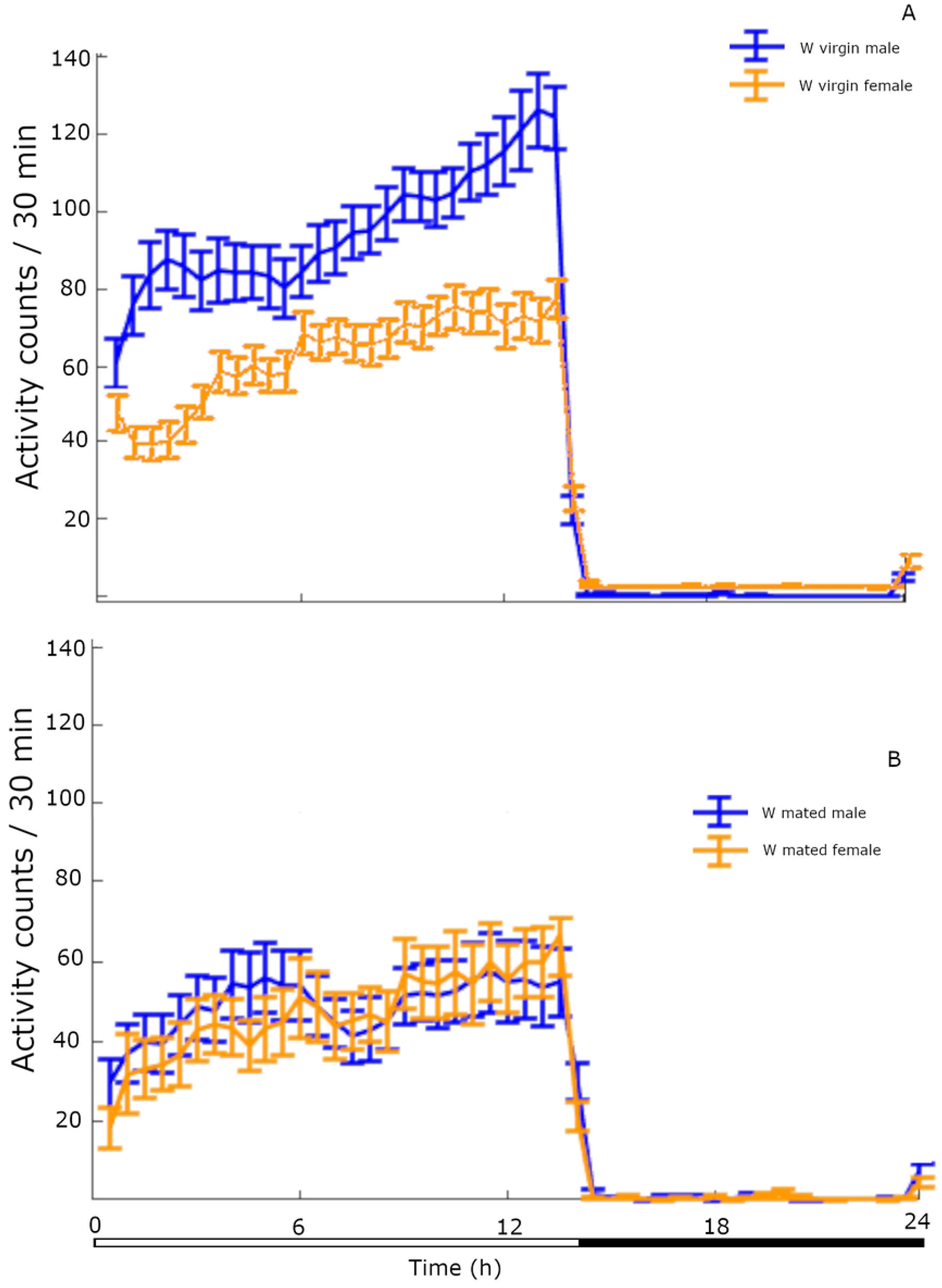
Locomotor activity levels (SE) of W virgin male and female flies (A) and W mated male and female flies (B) (*n* = 32 W virgin flies of each sex, *n* = 12 W mated males, *n* = 14 W mated females averaged across 5 days of monitoring).

The pattern of locomotor activity of W mated males and females during 24-h in 30 min bins is shown in Fig 1B. The mean locomotor activity level (SE) of W mated males and W mated females during the LP was 1153.1 (174.0) counts and 1081.9 (181.5) counts respectively and did not differ significantly (2-tailed *t*-test = .283, *df* = 24, *P* = .779). The mean locomotor activity level (SE) of W mated males and W mated females during the DP was 208.2 (30.5) counts and 219.0 (26.3) counts respectively and did not differ significantly (2-tailed *t*-test = −0.266, *df* = 23, *P* = .792).

As shown in Fig 2, mating affects the locomotor activity of males but not of females. The mean locomotor activity was significantly higher in virgin W males compared to mated ones during LP (2-tailed *t*-test = 4.020, *df* = 24, *P* = .0005) but not during DP (2-tailed *t*-test = 1.383, *df* = 19, *P* = .182). However, virgin and mated W females did not differ significantly in their mean locomotor activity during LP (2-tailed *t*-test = 1.553, *df* = 19, *P* = .136) or DP (2-tailed *t*-test = .889, *df* = 23, *P* = .382).

**Fig 2.**
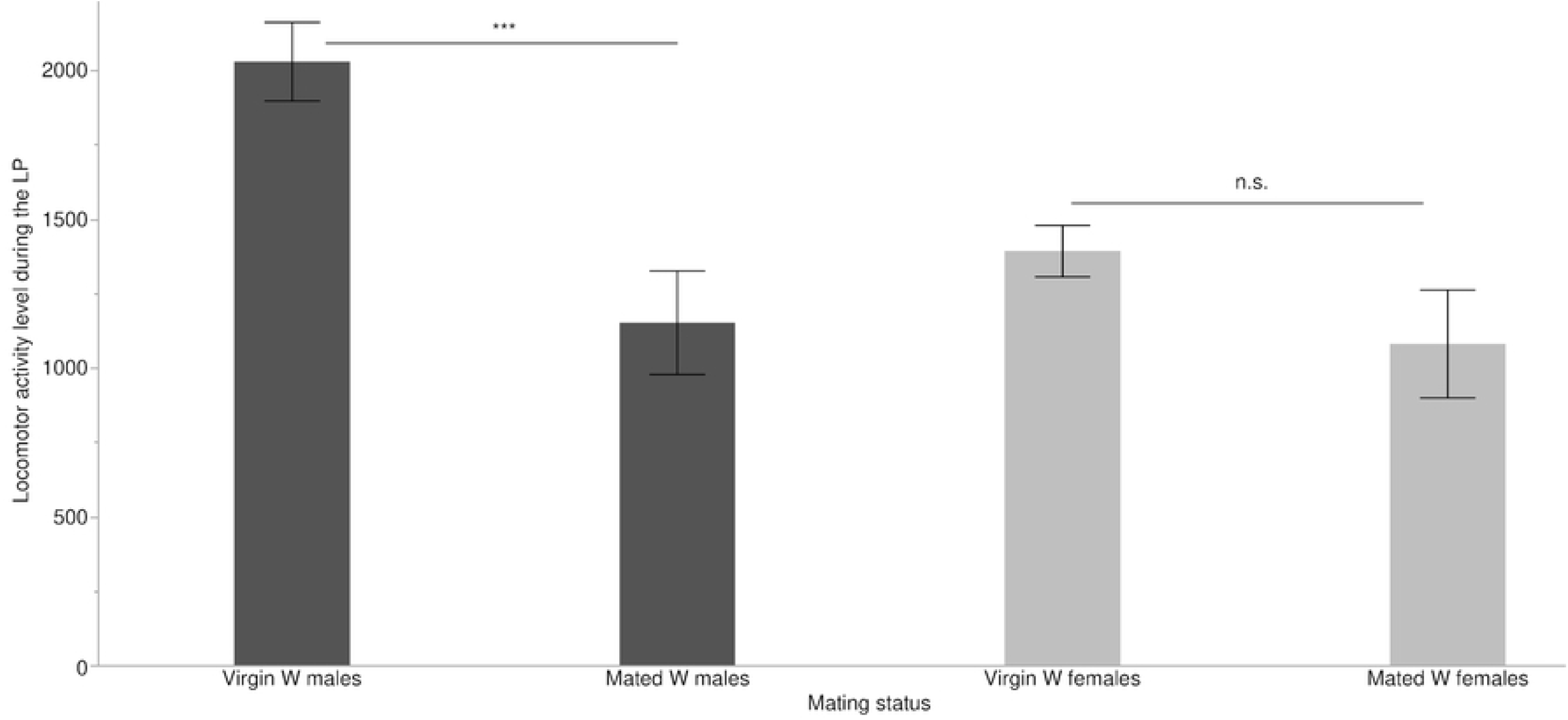
Mean (SE) level of locomotor activity of W virgin and mated flies of both sexes during the LP (*n* = 32 W virgin flies of each sex, *n* = 12 W mated males and *n* = 14 W mated females, averaged across 5 days of monitoring) (*** *P <* .*001*).

#### 3.1.2. Locomotor activity of AR flies

The pattern of locomotor activity of AR virgin males and females in 30-min bins across 24 h is shown in Fig 3. The mean locomotor activity level (SE) of AR virgin males and AR virgin females during the LP was 363.0 (63.2) counts and 324.3 (45.6) counts respectively. Contrary to the W flies, in AR flies there was no statistical difference between females and males during the LP (2-tailed *t*-test = .467, *df =* 27, *P* = .623). The mean locomotor activity level (SE) of AR virgin males and AR virgin females during the DP was 100.5 (10.5) counts and 68.6 (9.6) counts respectively. In contrast to the W flies, there was a statistical difference in locomotor activity between the AR female and male flies during the DP (2-tailed *t*-test = 2.236, *df* = 29, *P* = .0329).

**Fig 3.**
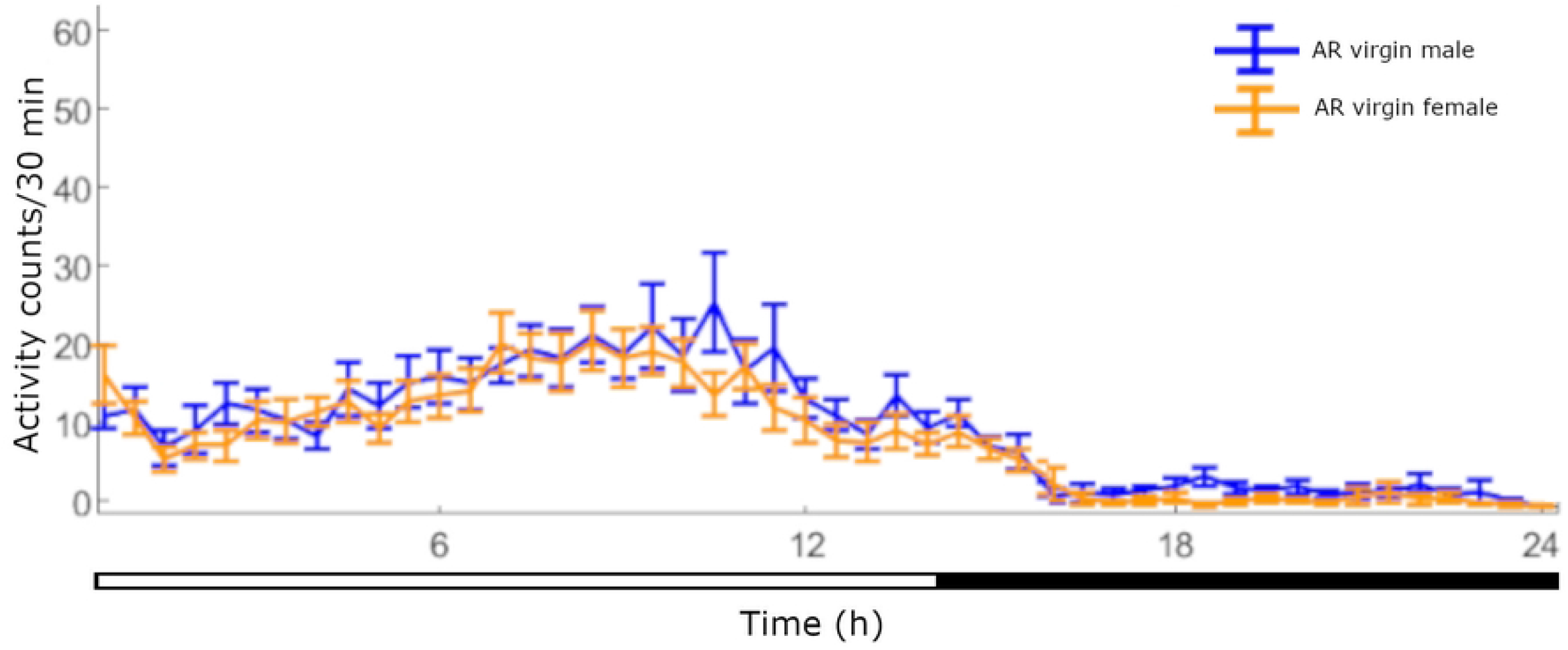
Locomotor activity levels (SE) of AR virgin male and female flies (*n* = 16 flies of each sex, averaged across 5 days of monitoring).

#### 3.1.3. Comparison of locomotor activity between W and AR virgin flies

There was significant difference in the mean locomotor activity during the LP between W and AR virgin flies of both sexes (2-tailed *t*-test = 11.398, *df* = 42, *P* < .0001 for males and *t*-test = 10.926, *df* = 43, *P* < .0001 for females) (Fig 4).

**Fig 4.**
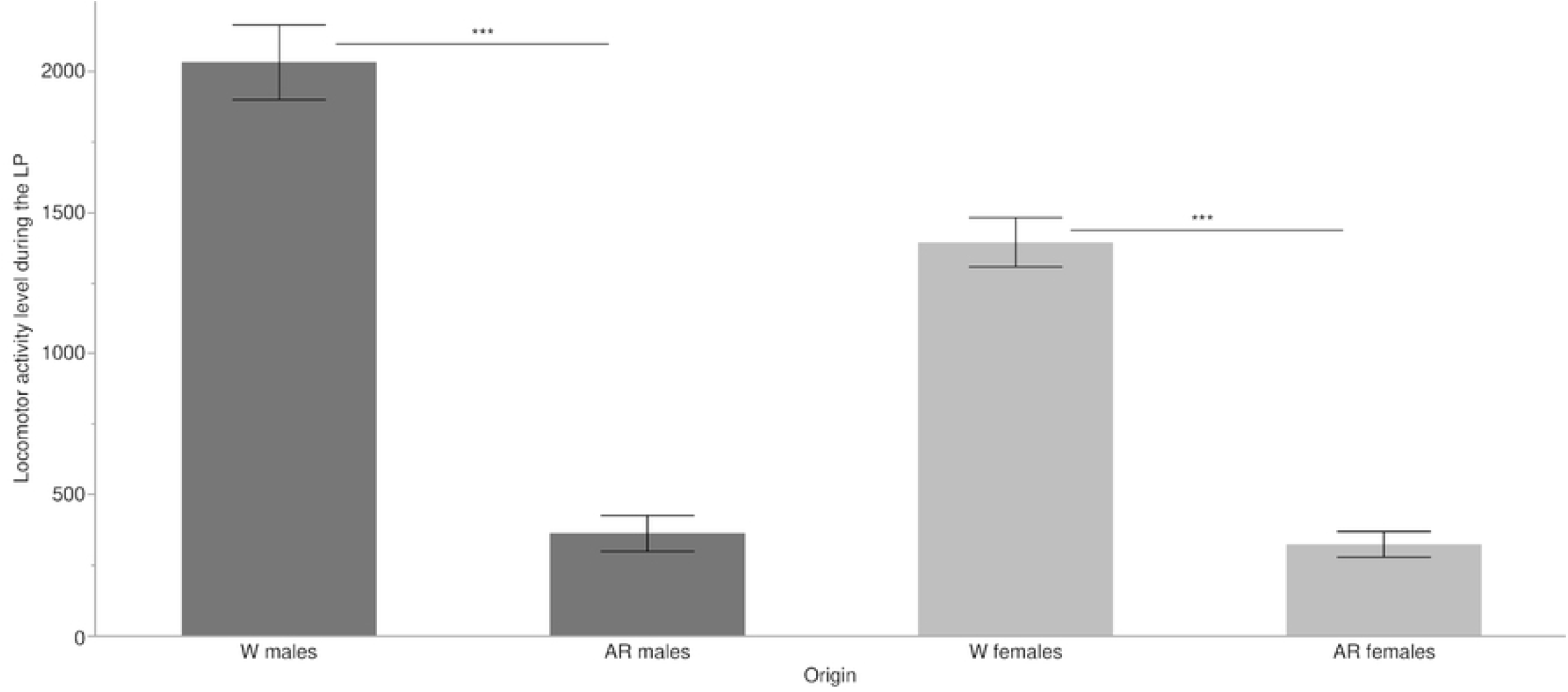
Mean (SE) levels of locomotor activity of W virgin and AR virgin flies as total counts of activity during the LP (*n* = 32 W flies of each sex, *n* = 16 AR flies of each sex, averaged across 5 days of monitoring) (*** *P <* .*001*).

### 3.2. Rest and sleep episodes

#### 3.2.1. Rest and sleep episodes of W flies

During the LP, the mean number of rest episodes (SE) of W virgin males was 12.1 (1.1) with mean duration (SE) 15.7 (1.5) min. The mean number of rest episodes (SE) of W virgin females was 16.8 (1.2) with mean duration (SE) 15.5 (0.8) min. There was significant difference between virgin male and female flies in the mean number of rest episodes (2-tailed *t*-test = −2.697, *df* = 61, *P* = .009) but not in their mean duration (2-tailed *t*-test = .193, *df* = 45, *P* = .847). During the DP, the mean number of sleep episodes (SE) of W virgin males was 2.9 (0.2) with mean duration (SE) 309.0 (21.1) min. The mean number of sleep episodes (SE) of W virgin females was 3.8 (0.2) with mean duration (SE) 255.8 (16.8) min. There was significant difference between the two sexes in the mean number of sleep episodes (2-tailed *t*-test = −2.602, *df* = 58, *P* = .011) but not in their mean duration (2-tailed *t*-test = 1.708, *df* = 58, *P* = .0902).

During the LP, the mean number of rest episodes (SE) of W mated males was 16.0 (2.0) with mean duration (SE) 77.5 (22.3) min. The mean number of rest episodes (SE) of W mated females was 19.6 (1.8) with mean duration (SE) 47.9 (13.3) min. There was no significant difference between the two sexes in the mean number of rest episodes (2-tailed *t*-test = −1.261, *df* = 28, *P* = .217) or their mean duration (2-tailed *t*-test = 1.133, *df* = 23, *P* = .268). During the DP, the mean number of sleep episodes (SE) of W mated males was 4.0 (0.5) with mean duration (SE) 250.2 (39.9) min. The mean number of sleep episodes (SE) of W mated females was 4.3 (0.3) with mean duration (SE) 219.2 (29.6) min. There was no significant difference between the two sexes in mean number of sleep episodes (2-tailed *t*-test = −0.502, *df* = 26, *P* = .619) or their mean duration (2-tailed *t*-test = .624, *df* = 26, *P* = .537).

W mated males differed in the mean duration of rest episodes compared to W virgin males (2-tailed *t*-test = −2.747, *df* = 14, *P* = .0156), but no difference was detected in the number of rest episodes or the number and mean duration of sleep episodes between the two groups. Similarly, W virgin and mated females differed in the mean duration of rest episodes (2-tailed *t*-test = −2.426, *df* = 15, *P* = .0282), but not in their number nor in the mean duration and number of sleep episodes (Figs 5 and 6).

**Fig 5.**
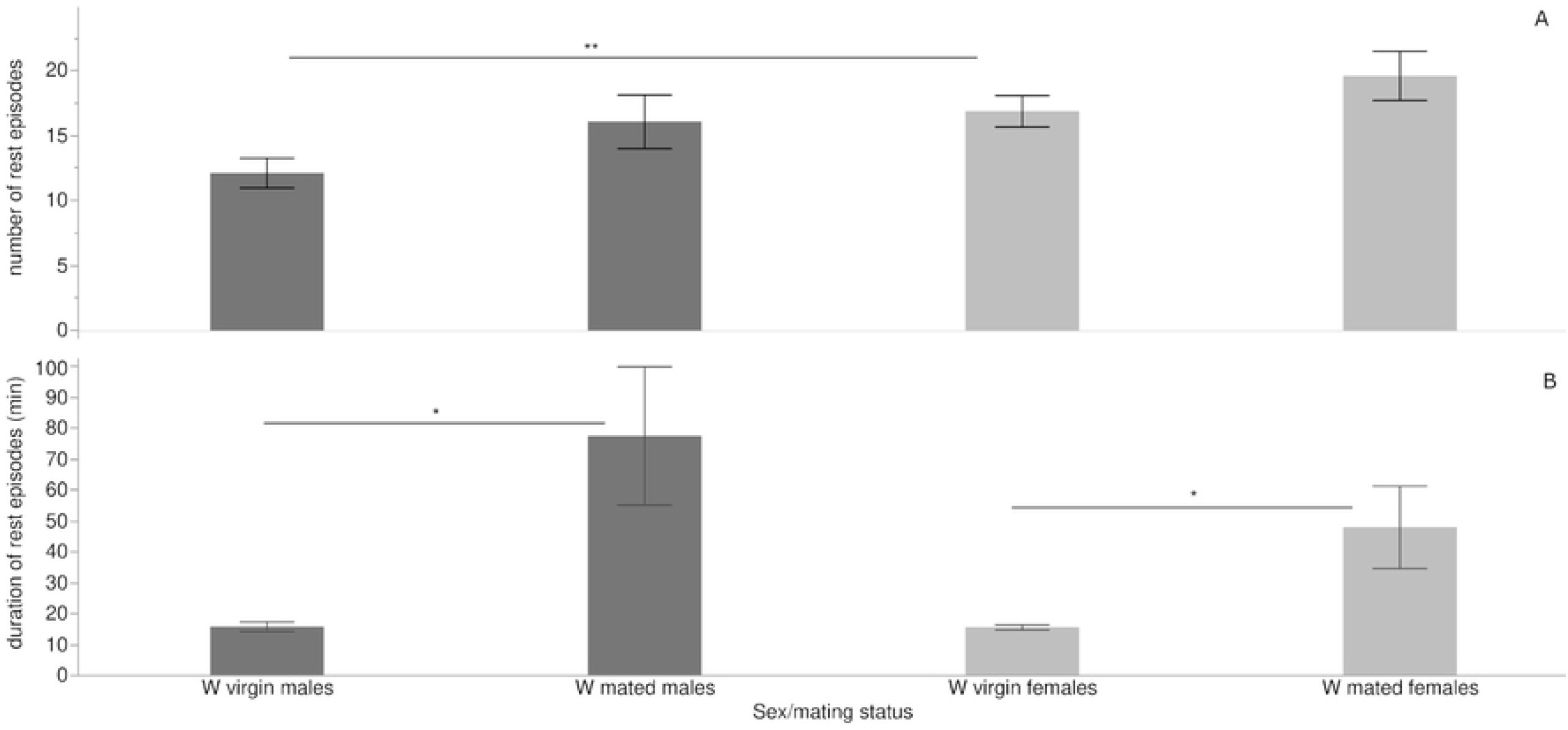
Mean (SE) number (A) and duration (B) of rest episodes of W virgin and mated flies of both sexes during the LP (* *P<* .*05*, ** *P <* .*01*).

**Fig 6.**
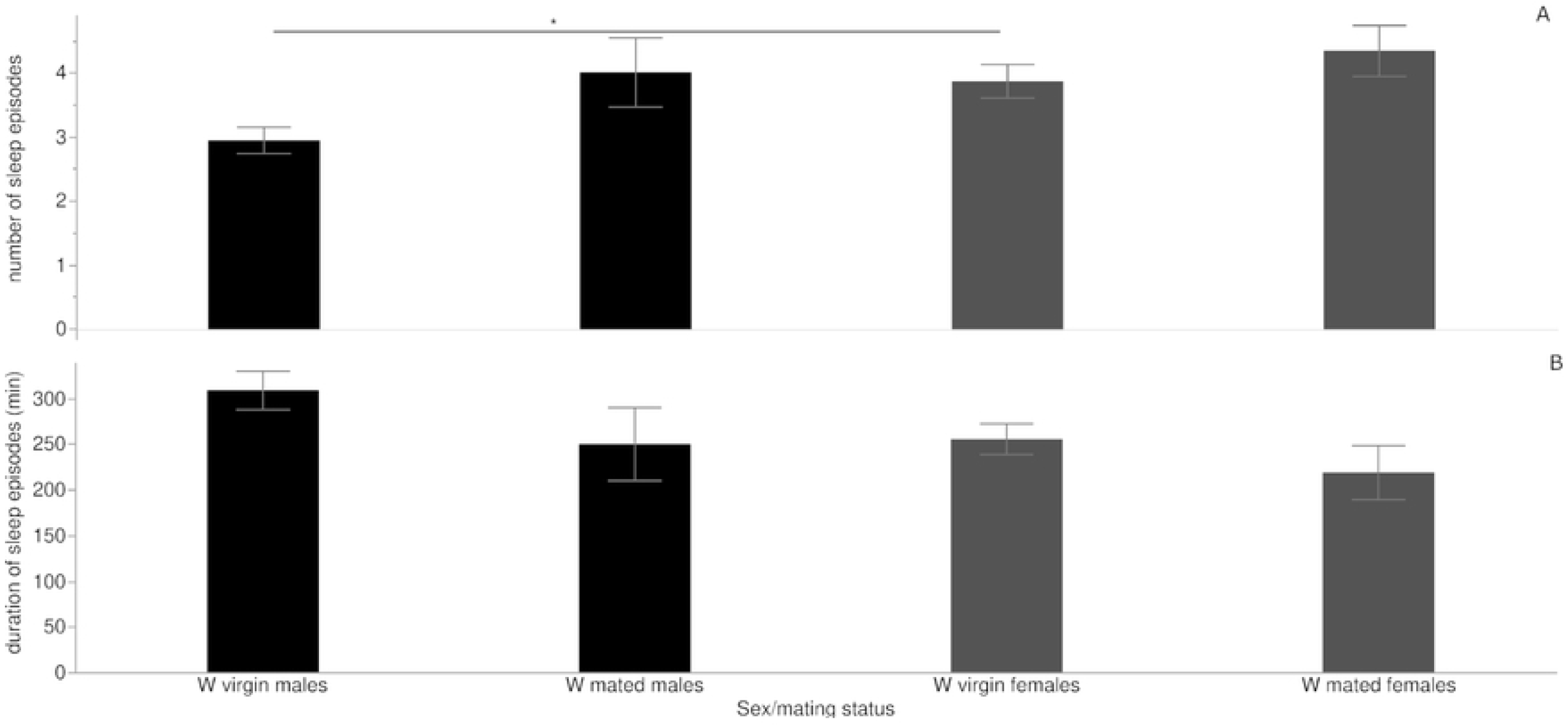
Mean (SE) number (A) and duration (B) of sleep episodes of W virgin and mated flies of both sexes during the DP (* *P<* .*05*).

#### 3.2.2. Rest and sleep episodes of AR flies

During the LP, the mean number of rest episodes (SE) of AR virgin males was 9.7 (0.9) with mean duration (SE) 110.6 (15.5) min. The mean number of rest episodes (SE) of AR virgin females was 16.0 (1.0) with mean duration (SE) 49.4 (5.0) min. There was significant difference between the two sexes in mean number of rest episodes (2-tailed *t*-test = −4.599, *df* = 29, *P* < .0001) and mean duration (2-tailed *t*-test = 3.735, *df* =18, *P* = .0015). During the DP, the mean number of sleep episodes (SE) of AR virgin males was 9.3 (0.7) with mean duration (SE) 116.3 (15.2) min. The mean number of sleep episodes (SE) of AR virgin females was 5.6 (0.4) with mean duration (SE) 194.1 (25.0) min. There was significant difference between the two sexes in number of sleep episodes (2-tailed *t*-test = 4.243, *df* = 24, *P* = .0003) and mean duration (2-tailed *t*-test = −2.656, *df* = 24, *P* = .0136).

#### 3.2.3. Comparison of rest/sleep episodes between W and AR male flies

W virgin males and AR virgin males differed in the mean number of sleep episodes (2-tailed *t*-test = −8.142, *df* = 17, *P* < .0001), their mean duration (2-tailed *t*-test = 7.512, *df* = 45, *P* < .0001) and also in the mean duration of rest episodes (2-tailed *t*-test = −6.866, *df* = 15, *P* < .0001), but not in their number (2-tailed *t*-test = 1.633, *df* = 44, *P* = .109). Interestingly, the mean number and duration of rest or sleep episodes for AR males, did not differ between the LP and DP (Fig 7).

**Fig 7.**
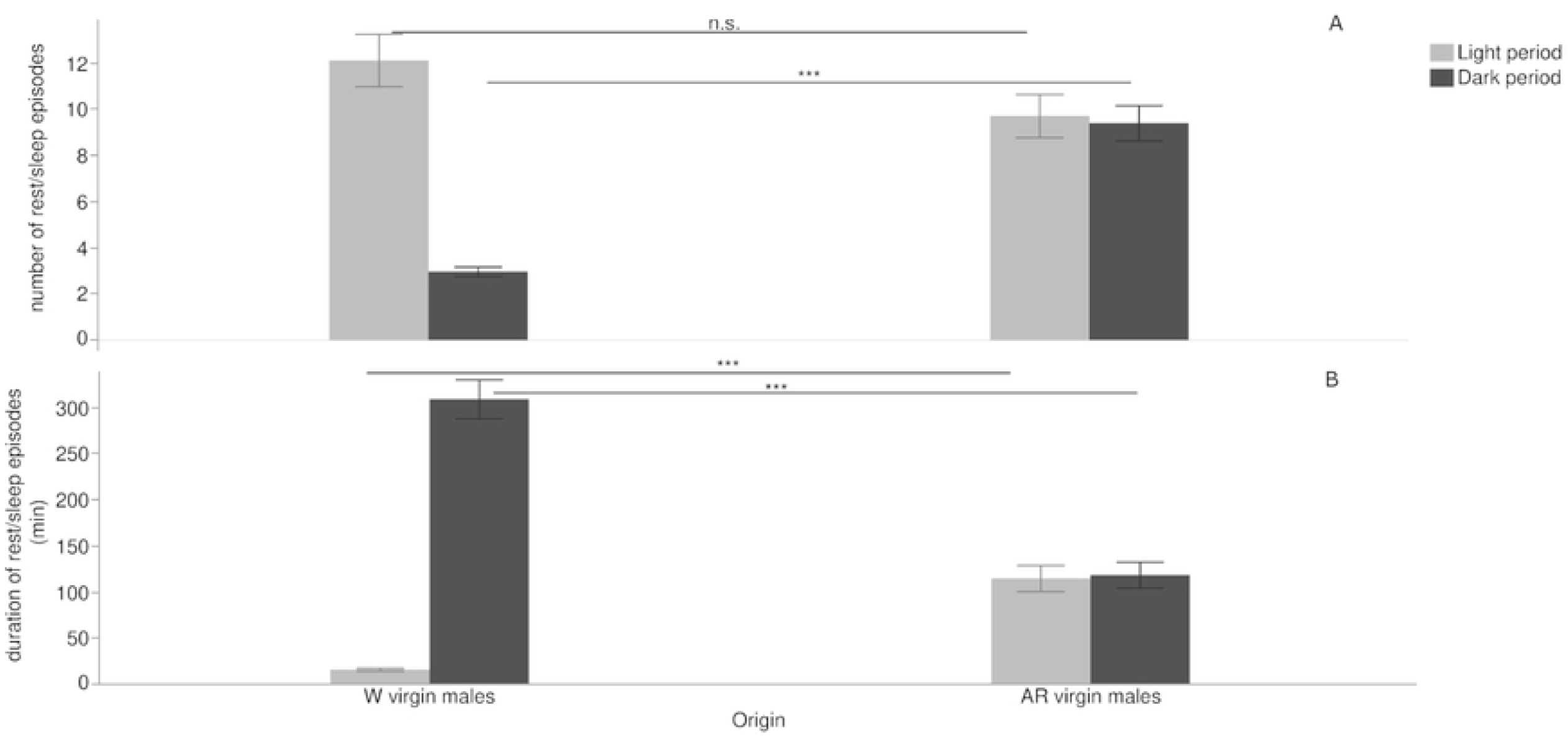
Mean (SE) number (A) and duration (B) of rest episodes (light grey color) and sleep episodes (dark grey color) for W and AR virgin males (*** *P<* .*001*).

## 4. Discussion

This study indicates that artificial rearing, mating status (virginity) and light and dark phases of photoperiod affect the locomotor activity of the olive fruit fly adults. W olive fruit flies are mostly active during the light period and bouts of inactivity (rest episodes) during this period have mean duration of 15 min. Mate searching and courtship in this species are done in late evening and W virgin males have exhibited increasing locomotor activity towards the end of the light period (Fig 1) in accordance to [34]. Locomotor activity of W flies during the dark period takes place mostly during the first hours after lights off. Inactivity bouts during the DP (sleep episodes) have a mean duration of 255-300 min for females and males respectively.

W females have lower locomotor levels than males, perhaps due to their heavier body and that they are monogamous. W mated males have reduced locomotor activity levels and rest episodes of longer duration compared to W virgin ones. However, W mated females have similar locomotor levels compared to W virgin ones, but have rest episodes of longer duration, which may be due to oviposition behavior. AR flies have lower locomotor activity levels compared to W ones, in accordance to many studies with mass reared insects [35]. The peak of the AR virgin males’ locomotor activity is earlier than the W virgin ones (Fig 5), and earlier mating times have been observed in laboratory adapted populations [36, 37]. Furthermore, we found that AR virgin males have higher number of sleep episodes of shorter mean duration compared to W virgin males.

The differences that have been detected in the level of activity and sleep patterns between W and AR male olive fruit flies can be attributed to laboratory adaptation [38] and could impact the AR flies’ survival, dispersion and ability to compete with the wild population. Increased locomotor activity has been associated with higher mating success in Tephritidae [39]. Mass reared *B. curcubitae* have reduced flight ability than the wild flies [40], and laboratory adaptation can change the biological traits of weevils [41], affect the fitness of parasitoids used for biological control [42], or lead to loss of stress resistance to *D. melanogaster* [43].

Bertolini et al. [5], have compared the locomotor activity between of a wild-type and a self-limiting strain of *B. oleae*, the wild-type strain refering to *Argov* and *Democritus* strains that are laboratory adapted. These strains’ day activity/h of male flies was found to be 17 counts/h and is similar with our findings of AR mature males’ day activity/h (363 counts/14 h = 25.9 counts/h). The reduced activity and high mortality of flies that Bertolini et al. noticed, could have been caused because the tubes used to house the flies in the LAM device had 10 mm diameter, while we used larger tubes (25 mm diameter), allowing the olive fruit flies to move more freely and be less stressed. Circadian clock regulates daily rhythms of animal physiology and behavior, which are entrained by environmental stimuli. Sleep is one of the established circadian behavior, which is also associated with health status. In Drosophila, genetic dissections show that sleep is regulated by the circadian clock and the circadian clock in the *B. oleae* has a *Drosophila*-like organization [5]. Mutations of the core clock genes have abnormal sleep, while the peptidergic clock neurons regulate arousal as well as sleep stability [44]. Laboratory-adapted *B*.*curcubitae* strains that during domestication process were artificially selected for short larval developmental time and early reproductive age exhibited changes in traits like shorter circadian periods and later time of mating in the day compared to the wild flies [45]. These changes could be attributed to artificial selection for clock genes that pleiotropically control circadian rhythm and the time of mating [46].

Our study showed that AR olive fruit flies had more fragmented night sleep and less total time of night sleep compared to W flies. Sleep bouts are shortened resulting in fragmented night sleep, with implications on their overall fitness and longevity and reduced night sleep quality might affect their day activity levels in humans [47]. When sleep is insufficient, important brain processes such as learning and memory are affected, and, when sleep is insufficient over the short term, this effect can be reversed by supplemental sleep [48].

Reduced levels of day activity for AR flies could be explained to poor night sleep quality. There are two processes, the circadian clock and the sleep homeostat that work together to regulate sleep. Circadian clock is responsible regulating the oscillation of sleep pressure during the day, while sleep homeostat conveys the need for sleep depending on the duration and quality of previous wake periods. If sleep is disrupted, the homeostat process is able to overcome the circadian process, and induce sleep into a period of the day that is normally dedicated to other activities, such as feeding and mating. This compensatory sleep is often referred to as “rebound sleep.” In insects, sleep also has an impact on fitness and can affect reproductive output and development [49]. Importantly, sleep-deprived males show reduced courtship behavior. Also, males lacking a functional clock show a significant decline in the quantity of sperm in *D. melanogaster*, for which it was demonstrated that clock genes are rhythmically and autonomously expressed in testes and seminal vesicles of male flies, suggesting that these tissues harbor a circadian system important for optimal sperm output and fertility [50].

Finally, the same experiments should be repeated in natural conditions, because the natural environment provides much richer cycling environmental stimuli than the laboratory and generally accepted adaptive and mechanistic explanations for fly circadian behavior from laboratory experiments may require some revision if they are to account for rhythmicity in nature [51].

